# Neural population dynamics underlying expected value computation

**DOI:** 10.1101/2020.06.01.127738

**Authors:** Hiroshi Yamada, Yuri Imaizumi, Masayuki Matsumoto

## Abstract

Computation of expected values, i.e., probability times magnitude, seems to be a dynamic integrative process performed in the brain for efficient economic behavior. However, neural dynamics underlying this computation remain largely unknown. We examined (1) whether four core reward-related regions detect and integrate the probability and magnitude cued by numerical symbols and (2) whether these regions have distinct dynamics in the integrative process. Extractions of mechanistic structure of neural population signal demonstrated that expected-value signals simultaneously arose in central part of orbitofrontal cortex (cOFC, area 13m) and ventral striatum (VS). These expected-value signals were incredibly stable in contrast to weak and/or fluctuated signals in dorsal striatum and medial OFC. Notably, temporal dynamics of these stable expected-value signals were unambiguously distinct: sharp and gradual signal evolutions in cOFC and VS, respectively. These intimate dynamics suggest that cOFC and VS compute the expected-values with unique time constants, as distinct, partially overlapping processes.

## Introduction

Economic behavior requires a reliable perception of the world for maximizing benefit (Houthakker, 1950; Samuelson, 1950; Savage, 1954; Von Neumann & Morgenstern, 1944). Such maximization is achieved in the first place by computing expected values (i.e. probability times magnitude) (Glimcher et al., 2008) in the brain, which seems to be a dynamic process to detect and integrate the probability and magnitude to yield expected value signals. Indeed, humans and animals behave as if they compute the expected values (Glimcher et al., 2008; Kahneman & Tversky, 1979; Stephens & Krebs, 1986). One salient example, discovered over a century ago and repeatedly measured, is human economic behavior, in which a series of models originating from the standard theory of economics (Von Neumann & Morgenstern, 1944) has been developed to describe efficient economic behavior. Despite the ubiquity of this phenomenon, a dynamic integrative process to compute the expected values from probability and magnitude remain largely unknown.

Substantial previous researches in the past two decades have shown that many regions of the brain process rewards in terms of signaling the probability and/or magnitude, mostly during economic choice behavior (Barraclough et al., 2004; Eshel et al., 2016; Hirokawa et al., 2019; Lopatina et al., 2016; Ma & Jazayeri, 2014; Platt & Glimcher, 1999; Roesch et al., 2009; Rudebeck & Murray, 2014; Tobler et al., 2005; Xie & Padoa-Schioppa, 2016; Yamada et al., 2018; Zhou et al., 2019). In these previous literature, the computation of expected values are assumed to be processed by these regions without their neural dynamics, in line with an expected value theory shared across multiple disciplines (Glimcher et al., 2008; Stephens & Krebs, 1986; Sutton & Barto, 1998; Von Neumann & Morgenstern, 1944). Neuroimaging studies in humans and non-human primates have also suggested that multiple brain regions in the reward circuitry (Haber & Knutson, 2010) are involved in this computational process (Fouragnan et al., 2019; Howard et al., 2015; Howard & Kahnt, 2017; Hsu et al., 2009; Levy & Glimcher, 2012; J. O’Doherty et al., 2004; Papageorgiou et al., 2017; Tom et al., 2007), although the underlying neural mechanism has not been elucidated because of the limited time resolution of current neuroimaging techniques (Goense & Logothetis, 2008; Milham et al., 2018). These reward-related brain regions may employ the expected-value computation, however, none have captured and compared the temporal aspects of neural activities in the multiple candidate brain regions. These observations have led us to test the hypothesis that neural population dynamics within subsecond-order time resolutions (Chen & Stuphorn, 2015; Churchland et al., 2012; Mante et al., 2013; Murray et al., 2017; Takei et al., 2017) play key roles in the expected-value computation, that is the detection and integration of the probability and magnitude on multiple neural population ensembles.

We targeted reward-related cortical and subcortical structures of non-human primates (Haber & Knutson, 2010) including the central orbitofrontal cortex (cOFC, area 13M), the medial orbitofrontal cortex, (mOFC, area 14O), the dorsal striatum (DS, caudate nucleus), and ventral striatum (VS), all of which are known to represent the neural correlate with stimulus values during economic choice behavior. To dissociate the integrative process to compute expected values from a choice process employed during economic choices (Chen & Stuphorn, 2015; Gardner et al., 2019; Yoo & Hayden, 2020), we recorded the neural activity in a non-choice situation; monkeys perceive and compute the expected values from probability and magnitude symbols. We used a recently developing mathematical approach, called state space analysis (Chen & Stuphorn, 2015; Churchland et al., 2012; Mante et al., 2013; Murray et al., 2017), to test how the expected-values computation is processed within each of the four neural population ensembles in an order of 10^−2^-second time resolution. We found that the cOFC and VS neural populations coincidently detect and integrate the probability and magnitude information to yield the expected value signals on neural population assembles. These signals are maintained in a stable state, though the cOFC signal sharply develops in contrast to gradual evolution of the VS signal. Contrastingly, the signals in the mOFC and DS were weak and/or fluctuated. Thus, we conclude that the cOFC and VS neural populations employ a common integrative computation of expected values from probability and magnitude.

## Results

### Task and behavior in monkeys

Based on the vast literature on human behavioral economics and by harnessing the well-developed visual and cognitive abilities in non-human primates, we designed a behavioral task in which monkeys estimated the expected values of rewards from numerical symbols, mimicking events performed by humans. The task involved a visual pie-chart that included two numerical symbols associated with the probability and magnitude of fluid rewards with great precision. After monkeys fixated the central grey target, a visual pie-chart comprising green and blue pies was presented (Figure 1A). The number of the green pies indicated the magnitude of fluid rewards in 0.1 mL increments (0.1-1.0 mL). Simultaneously, the number of the blue pies indicated the probability of reward in 0.1 increments (0.1-1 where 1 indicates a 100% chance). After a 2.5 second delay, the visual pie-chart disappeared, and a reward outcome was provided to the monkeys with the indicated amount and probability of reward, unless no reward was given. Under this experimental condition, the expected values of rewards are defined as the probability multiplied by the magnitude cued by the numerical symbols (Table. S1).

**Figure 1.**
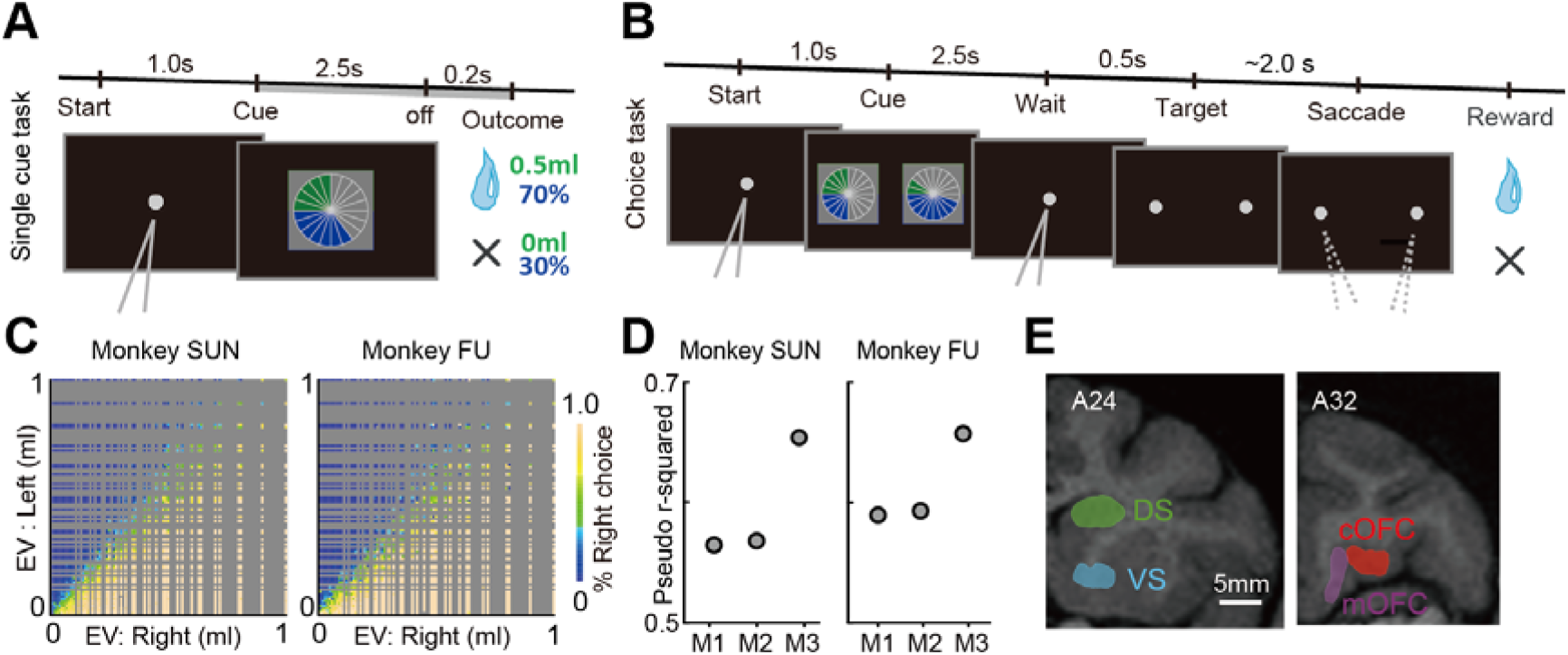
Task and behavior for recording neural activity. (**A**) Sequence of events during the single cue task. A single visual pie-chart comprising green and blue pies was presented to the monkeys. (**B**) Choice task. Two visually displayed pie-charts were presented to the monkeys left and right to the center. After visual fixation on the re-appeared central target, the central fixation target disappeared, and monkeys chose either one of the targets by fixating on the target. A block of choice trials was interleaved between single cue trial blocks. During the choice trials, neural activity was not recorded. (**C**) Percentages of right target choice during the choice task plotted against the expected values of left and right target options. (**D**) Pseudo r-squared estimated in the three behavioral models. M1: number of pies. M2: probability and magnitude. M3: the expected values. (**E**) An illustration of neural recording areas based on sagittal magnetic resonance images. Neurons were recorded from medial (mOFC, 14O, orbital part of area 14) and central parts of the orbitofrontal cortex (cOFC, 13M, medial part of area 13) at A31-A34 anterior-posterior (A-P) level. Neurons were also recorded from the dorsal (DS) and ventral striatum (VS) at A21-A27. The white scale bar indicates 5 mm.

To examine whether the monkeys accurately perceived the expected values from the numerical symbols for probability and magnitude, we applied a choice task to the monkeys (Figure 1B). The two monkeys exhibited a near-efficient performance in selecting a larger expected value option among two alternatives during choice trials (Figure 1C). We examined which of the following three behavioral models best described the monkey’s behavior: model 1 (M1), monkeys make choices based on the number of pies; model 2 (M2), monkeys make choices based on the probability and magnitude; and model 3 (M3), monkeys make choices based on the expected values (Table. S1). Comparisons of the model performances based on Akaike’s Information Criterion (AIC) and Bayesian Information Criterion (BIC) (Burnham & Anderson, 2004) revealed that the model 3 best explained the monkey’s behavior as indicated by the smallest AIC and BIC values (Figure S1). The model 3 consistently showed the highest pseudo r-squared values in each monkey (Figure 1D). These results indicated that the monkeys utilize the expected values estimated from the numerical symbols for probability and magnitude.

### Neural population data

We constructed four pseudo-populations of neurons by recording single-neuron activity during the single cue task (Figure 1A) from the DS (194 neurons), VS (144 neurons), cOFC (190 neurons) and mOFC (158 neurons) (Figure 1E). The four constructed neural populations exhibited changes in their activities at different times in the task trials (Figure S2A). Approximately 40 to 50 % of neurons in the four neural populations demonstrated peak activity during the cue presentation (Figure S2B, Chi-squared test, n = 686, *P* = 0.32, X^2^ = 3.55, df = 3), with several basic firing properties (Figure S2C-F). Strong peak activities with short-latency were observed in the cOFC (Kruskal-Wallis test, Latency: Figure S2C, n = 314, *P* = 0.013, H = 10.9, df = 3, Peak firing rate: Figure S2D, n = 314, *P* < 0.001, H = 32.1, df = 3). Activity changes were slow in the mOFC (Figure S2E, Kruskal-Wallis test, n = 314, *P* = 0.003, H = 13.4, df = 3). Baseline firing rates were the highest in the cOFC (Figure S2F, Kruskal-Wallis test, n = 686, *P* < 0.001, H = 60.3, df = 3). In short, the strong activity with short-latency frequently occurred in the cOFC in contrast to the phasic activity at various latencies in the DS and VS and relatively tonic and gradual activity changes in the mOFC.

### Conventional analyses for detecting the expected-value signals

We first applied common conventional analyses (linear regression, model selection based on AIC, and model selection based on BIC) to the four neural populations to examine neural modulations by the probability, magnitude, and expected values at a single neuron level (see Methods). During a fixed one second time window after the cue onset, these analyses showed that neurons in all the four brain regions signal the probability, magnitude, and expected values to some extent (Figure 2). For example, neurons signaling the expected values were found in each brain region (Figure 2A-H, Neural modulation pattern was defined as the Expected value type based on all the three analyses. Regression coefficients for probability and magnitude, A-B, DS, 6.17, P < 0.001 and 2.54, P = 0.007, C-D, VS: 7.14, P < 0.001 and 6.71, P < 0.001, E-F, cOFC, 8.55, P < 0.001 and 11.1, P < 0.001), while the classification of neural modulation types was dependent on the methods in some neurons (Regression coefficients for probability and magnitude, G-H, mOFC, 1.76, P = 0.032, and 0.50, P = 0.54, AIC-based model selection: Expected value type, linear regression: Probability type, and BIC-based model selection: non-modulated type). In addition, neurons signaling probability or magnitude were also observed in each brain region (Figure 2I-L, blue and green). Moreover, a subset of neurons in the cOFC and VS signaled high-risk high-return or low-risk low-return (Figure S3). These neurons were characterized as a strong activity elicited when the cue indicated a low probability and large magnitude (hence, high-risk high-return, Figure 2J and K, brown). Indeed, each neural population was composed of a mixture of these signals (Figure 2I-L), indicating that signals for the expected values and its components (i.e., probability and magnitude) appeared in each neural population during one second after the cue onset. If we analyzed these neural modulation patterns through a task trial (Figure 2M-P), there was no significant difference in the proportions of neural modulation types, except the VS (chi-squared test: DS, n = 104, df = 75, X^2^ = 91.4, *P* = 0.096; VS, n = 104, df = 75, X^2^ = 98.2, *P* = 0.037; cOFC, n = 104, df = 75, X^2^ = 83.2, *P* = 0.242; mOFC, n = 104, df = 75, X^2^ = 79.0, *P* = 0.353), indicating that these conventional analyses did not clearly provide evidence of temporal dynamics in the modulation patterns of neural populations. Thus, we developed an analytic tool to examine how the detection and integration of probability and magnitude are processed within these neural population ensembles.

**Figure 2.**
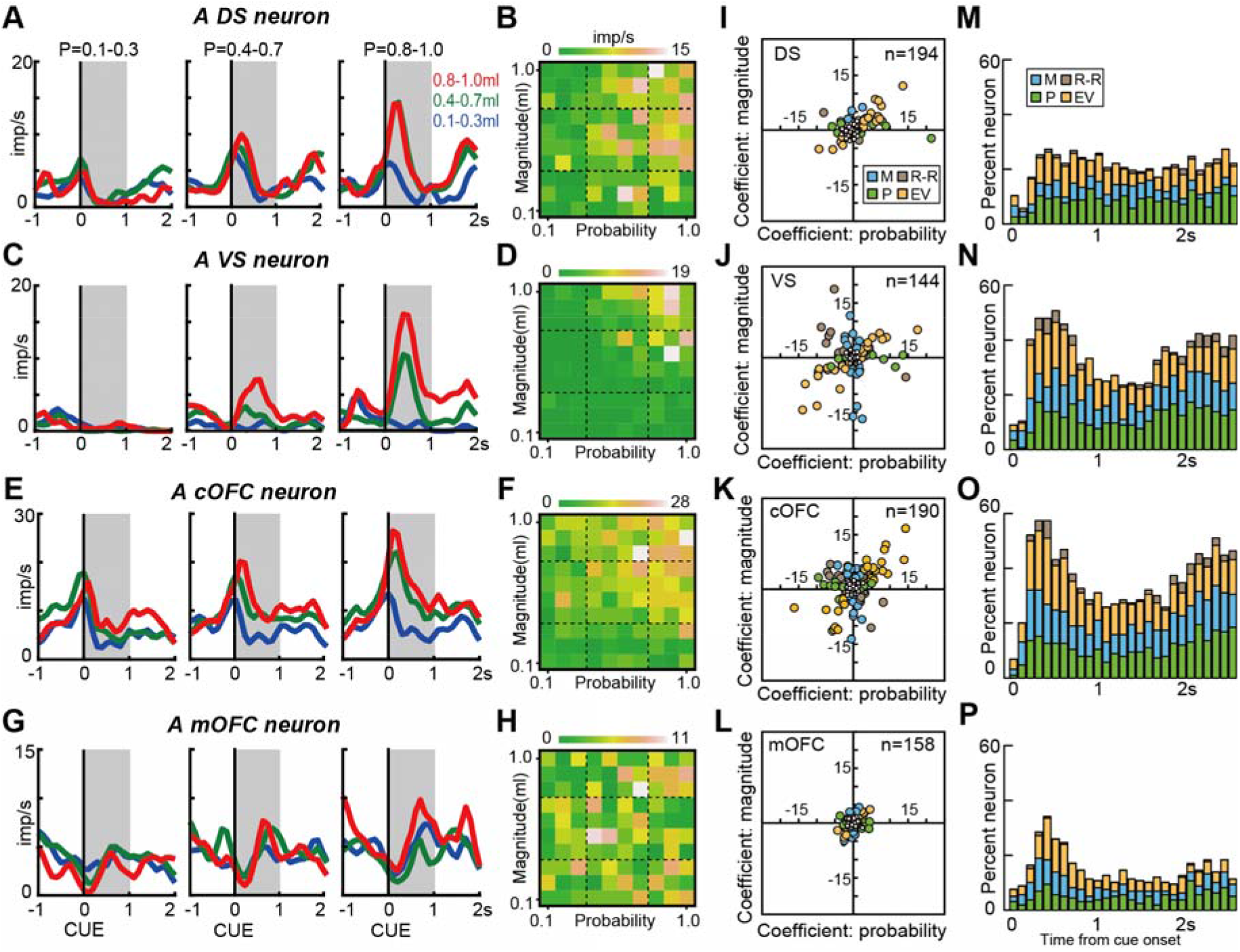
Expected-value signals detected by conventional analyses. (**A**) An example activity histogram of a DS neuron modulated by the expected values during the single cue task. The activity aligned to the cue onset is represented for three different levels of probability (0.1-0.3, 0.4-0.7, 0.8-1.0) and magnitude (0.1-0.3 mL, 0.4-0.7 mL, 0.8-1.0 mL) of rewards. Gray hatched time windows indicate a 1 s time window used to estimate the neural firing rates shown in **B**. (**B**) An activity plot of the DS neuron during the 1 s time window shown in **A** against the probability and magnitude of rewards. (**C-D**) same as **A-B**, but for a VS neuron. (**E-F**) same as **A-B**, but for a cOFC neuron. (**G-H**) same as **A-B**, but for a mOFC neuron. (**I-L**) Plots of the regression coefficients for the probability and magnitude of rewards estimated for all neurons in the DS (**I**), VS (**J**), cOFC (**K**), and mOFC (**L**). Filled colors indicate the neural modulation pattern classified by the BIC-based model selection. P: Probability type, M: Magnitude type, EV: Expected value type, and R-R: Risk-Return type. The non-modulated type is indicated by the small open circle. (**M-P**) Percentages of neural modulation types defined based on the BIC-based model selection through the cue presentation in the DS (**M**), VS (**N**), cOFC (**O**), and mOFC (**P**). The results using 0.1 s time window (without overlap) were shown. The classification of neural modulation types was dependent on the methods, although an overall tendency for the differences in neural modulations among the neural populations was consistent among the three analyses.

### State space analysis for detecting neural population dynamics

The state space analysis can provide temporal dynamics of neural population signal related to cognitive and motor performances (Churchland et al., 2012; Mante et al., 2013). In our lottery task, such population dynamics can describe how the expected values are evolved within the neural population ensembles. To describe how each neural population detects and integrates the probability and magnitude into the expected values, we represented each neural population signal as a time-series of vectors in the space of probability and magnitude in two steps. First, we used a linear regression to project a time series of each neural activity into the regression subspace composed of the probability and magnitude in each neural population. This step captures the across-trial variance caused by probability and magnitude moment-by-moment at a population level. Second, we applied a principal component analysis (PCA) to the time series of neural activities in the regression subspace in each neural population. This step determines the main feature of the neural population signal moment-by-moment in the space of probability and magnitude. Because activations are dynamic and changing over time, the analysis identified whether and how signal transformations occurred to convert the probability and magnitude into the expected values as a time-series of eigenvectors (Figure 3A). The directions of these eigenvectors capture the expected values as an angle moment-by-moment at a population level (Figure 3B). We evaluated properties of the eigenvectors for the first and second principal components (PC1 and PC2) in each neural population in terms of the vector angle, size, and deviance (Figure 3C). A stable population signal is described as a small variation in the eigenvector properties through a trial, whereas an unstable population signal is described as a large variation in the eigenvector’s properties. It must be noted that our procedure is a variant of the state space analysis in line with the use of linear regression to identify dimensions of a neural population signal (Chen & Stuphorn, 2015; Mante et al., 2013). It was not, however, aimed at projecting the population activity as trajectories in multi-dimensional space.

**Figure 3.**
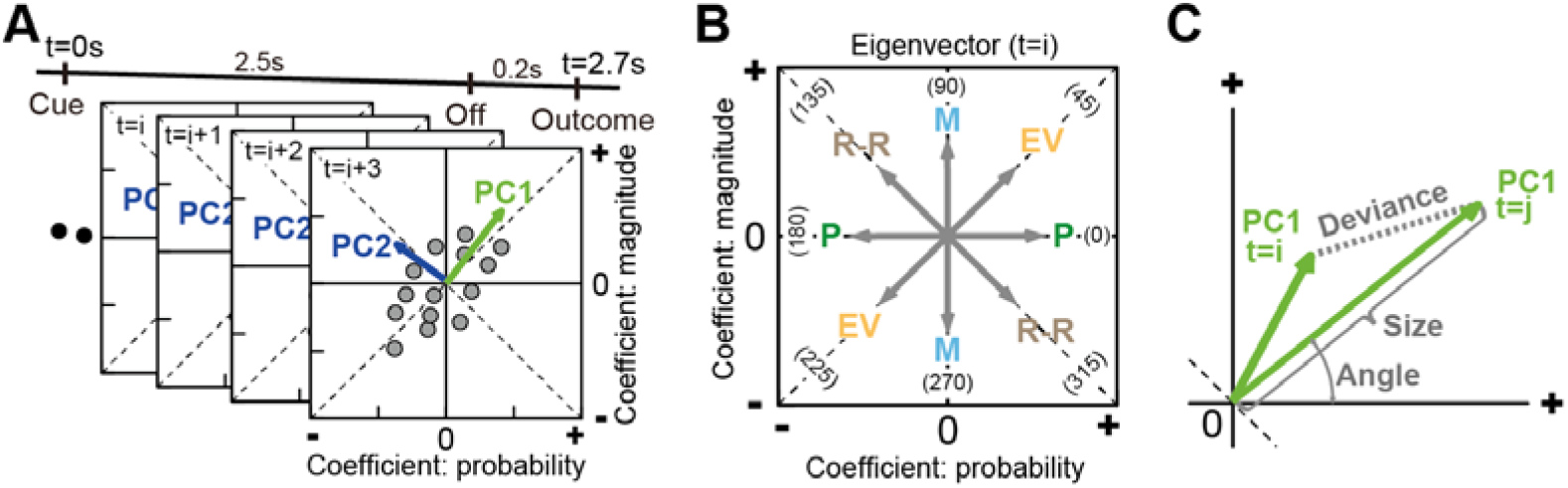
Schematic depictions for the analysis of neural population dynamics using PCA. (**A**) Time series of a neural population activity projected into a regression subspace composed of the probability and magnitude. A series of eigenvectors was obtained by applying PCA once to each of the four neural populations. PC1 and PC2 indicate the first and second principal components, respectively. The number of eigenvectors obtained by the PCA was 2.7 s divided by the analysis window size for the probability and magnitude; 27, 54, and 135 eigenvectors in 0.1, 0.05, or 0.02 s time window, respectively. (**B**) Examples of eigenvectors at time of *i* th analysis window for the probability and magnitude, whose direction indicates a signal characteristic at that time represented on the population ensemble activity. EV: expected values (45°, 225°), M: magnitude (90°, 270°), P: probability (0°,180°), R-R: risk-return (135°, 315°). (**C**) Characteristics of the eigenvectors evaluated quantitatively. Angle: vector angle from horizontal axis from 0° to 360°. Size: length of the eigenvector. Deviance: difference between vectors.

### Stable and unstable neural population signals

The eigenvector analyses yielded clear differences in neural population signals among the four populations (Figure 4A-D). We first confirmed adequate performance of the state space analysis indicated by the percentages of variance explained in each population (Figure 4A). The VS population exhibited the highest performance among the four neural populations, which was followed by the cOFC and DS populations, with the lowest performance exhibited by the mOFC population. Thus, performances to process the probability and magnitude information was distinct among the four neural populations.

**Figure 4.**
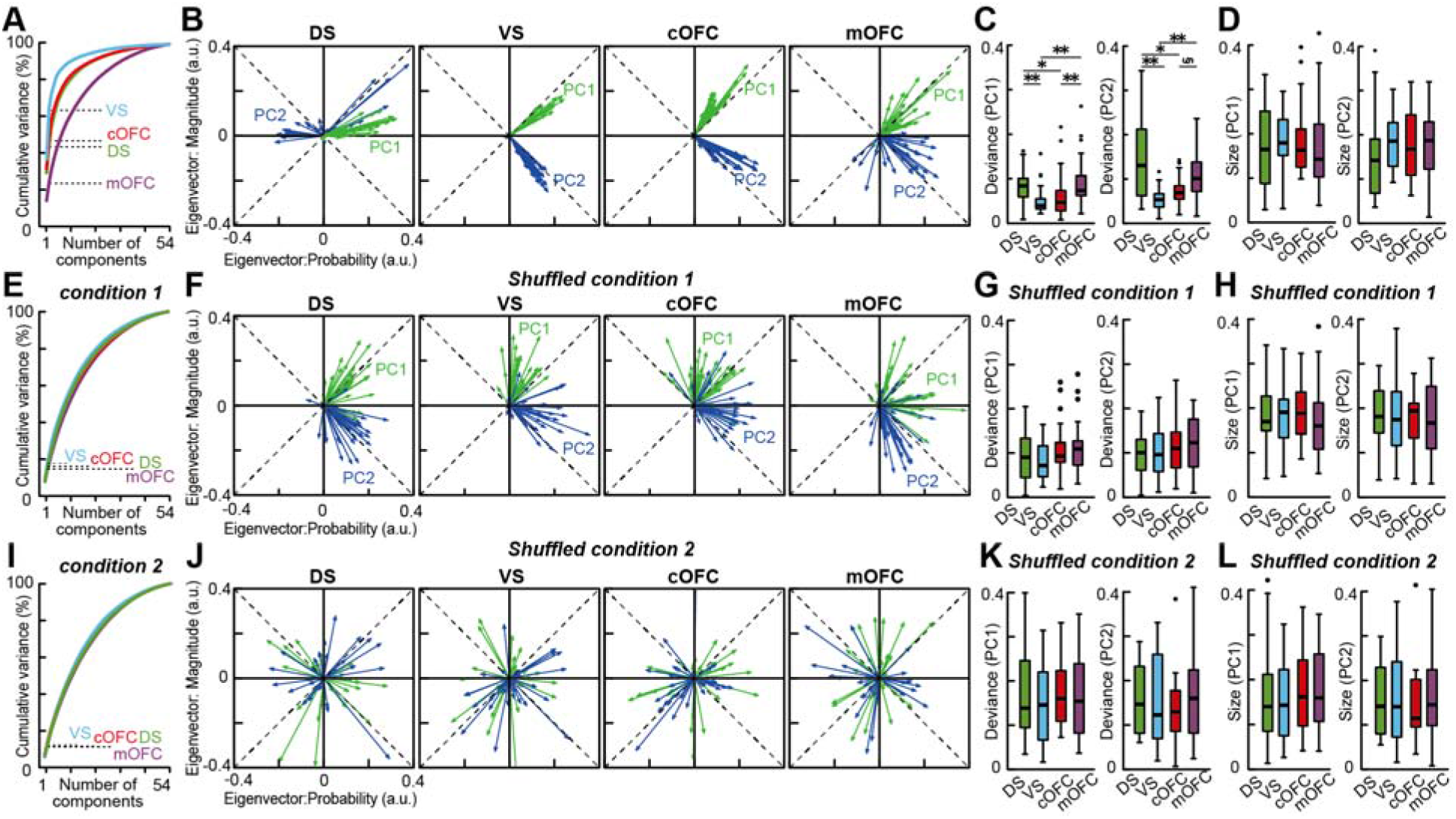
Neural populations provide stable expected-value signals in the VS and cOFC. (**A**) Cumulative variance explained by PCA in the four neural populations. Dashed line indicates percentages of variances explained by PC1 and PC2 in each neural population. (**B**) Overlay plots of the series of eigenvectors for PC1 and PC2 in the four neural populations. a.u. indicates arbitrary unit. (**C**) Box plots of the deviance from the mean vector estimated in each neural population for the PC1 (left) and PC2 (right). (**D**) Box plots of the vector size estimated in each neural population for the PC1 (left) and PC2 (right). (**E-H**) Same as **A-D**, but for the PCA under the shuffled condition 1. See Methods for details. (**I-L**) Same as **A-D**, but for the PCA under the shuffled condition 2. In **C-D**, **G-H**, and **K-L**, asterisks indicate statistical significance between two populations using Wilcoxon rank-sum test with Bonferroni correction for multiple comparisons (**,*, and § indicate statistical significance at *P* < 0.01, *P* < 0.05, and 0.05 < *P* < 0.06 (close to significance), respectively). The results are shown by using 0.1 s analysis window.

To characterize the whole structure of each neural population signal, we analyzed the aggregated properties of the eigenvectors without a temporal order of the vectors through a task trial. We first examined the properties of the eigenvectors for PC1. The aggregated eigenvectors revealed both stable and unstable neural population signals during the cue presentation (Figure 4B, green). The VS population exhibited the highest performance (37%) and the eigenvectors for PC1 were stable throughout cue presentation, with directions close to 45°, i.e., the expected value (Figure 4B, VS, vector angle, PC1, mean ± s.e.m., 37.5° ± 0.98, 7.5° difference from 45°). The cOFC population also exhibited a stable expected-value signal with the second-best performance (31%), but they were deviated more from the ideal expected-value signal (Figure 4B, cOFC, vector angle, PC1, mean ± s.e.m., 59.4° ± 1.16, 14.4° difference from 45°, Wilcoxon rank-sum test, n = 52, W = 122, *P* < 0.001). Stability of the vectors was the best in the VS and cOFC, as indicated by the smallest deviation from its mean vector among the four neural populations (Figure 4C, left, PC1). Thus, the VS and cOFC populations signaled the expected values in a stable manner.

In contrast, unstable population signals were observed in the DS and mOFC (Figure 4B, green). The DS population showed a considerable variability in their eigenvectors (Figure 4C, left, PC1) compared to those in the VS and cOFC neural populations. The signal carried by the DS neural population was close to 0°, i.e., the probability (Figure 4B, DS, vector angle, PC1, mean ± s.e.m., DS, 11.4° ± 1.72) with a performance closer to that of the cOFC (29%). The mOFC population exhibited a large variability in the eigenvectors (Figure 4B, mOFC, PC1, vector angle, mean ± s.e.m., 38.1° ± 5.80, Figure 4C, left, PC1) due to the poorest performance of PCA (14%), indicating a weak and fluctuating population signal. Thus, neural populations in the DS and mOFC did not signal the expected values due to the dynamic changes and weakness of the signals, respectively.

Second, we examined the properties of the eigenvectors for PC2. The eigenvectors for PC2 revealed another feature of the neural population signals, that reflected risk-return in the VS and cOFC (Figure S3, Figure 4B, blue, vector angle, PC2, mean ± s.e.m., VS, 306.7° ± 1.07, 8.3° difference from 315°, cOFC, 322.4° ± 1.94, 7.4° difference from 315°). The deviations from the ideal risk-return signal were not significantly different between the VS and cOFC populations (Wilcoxon rank-sum test, n = 52, W = 319, *P* = 0.737). These signals were equally stable in the VS and cOFC (Figure 4C, right, PC2). In clear contrast, the DS and mOFC signals were unstable and fluctuated more (Figure 4C, right, vector angle, PC2, mean ± s.e.m., DS, 64.8 ± 19.0, mOFC, 320.2 ± 8.77), similar to those observed for PC1 (Figure 4C, left, PC1). Thus, the VS and cOFC were key brain regions to signal risk-return as well as the expected values within their neural population ensembles, suggesting that integrated information of the probability and magnitude could be signaled in these neural populations.

To further examine the significance of these findings, we used a shuffle control procedure in two ways (see Methods). First, we randomly shuffled the allocation of probability and magnitude conditions to neural activity in each trial in each neuron (shuffled condition 1). If we shuffled the linear projection of neural activity into the regression subspace by this way, the neural population structure disappeared in all the four brain regions (Figure 4F). The performance of PCA for PC1 and PC2 were all below 20% (Figure 4E) and were significantly reduced from the observed data in all the four brain regions, even in the mOFC (Figure 5A, variance explained, *P* < 0.001 for all populations in PC1 and PC2). The deviance of eigenvectors under the shuffle control clearly increased in the cOFC and VS, which resulted in no significant differences among the four neural populations (Figure 4G, Kruskal-Wallis test, PC1, n = 104, df = 3, H = 4.99, *P* = 0.18, PC2, n = 104, df = 3, H = 2.78, *P* = 0.43). We also tested another shuffle control, in which the trial conditions were shuffled in each analysis window throughout a trial (shuffled condition 2). Under this full shuffled control, the PCA performances decreased further, albeit slightly (Figure 4I and Figure 5B), without significant differences among the four populations (Figure 4J-K, Deviance, Kruskal-Wallis test, PC1, n = 104, df = 3, H = 1.38, *P* = 0.71, PC2, n = 104, df = 3, H = 0.53, *P* = 0.91). Thus, these shuffle procedures appropriately evaluated the significance of our population findings.

**Figure 5.**
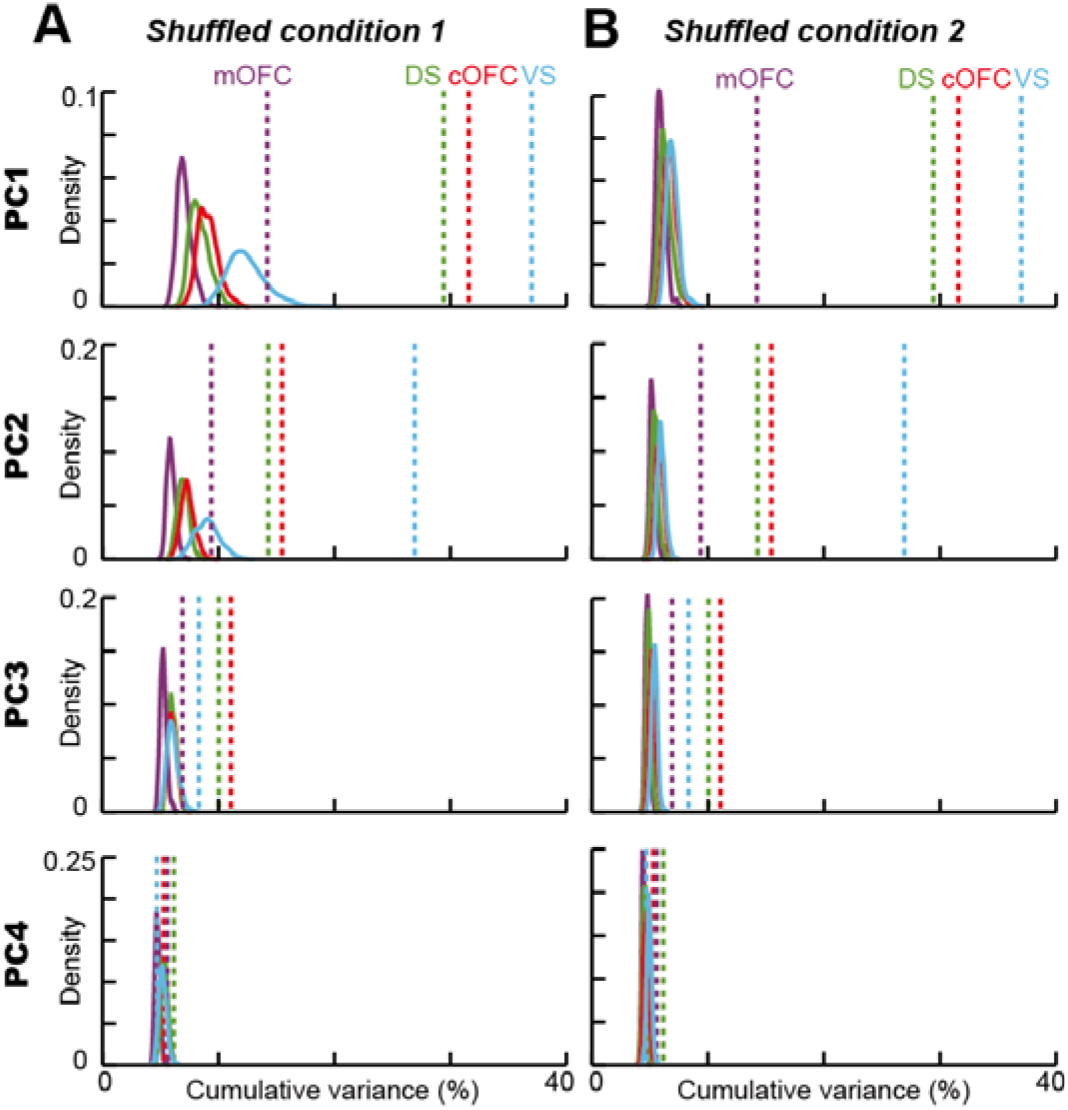
Probability density of the explained variances by PCA in the shuffled controls. (**A**) Probability density of variances explained by PCA for PC1 to PC4 under the shuffled condition 1 (see Methods for details). The probability density was estimated with 1,000 repeats of the shuffles in each neural population. (**B**) Probability density of variance explained by the PCA for PC1 to PC4 under the shuffled condition 2 (see Methods for details). The probability density was estimated with 1,000 repeats of the shuffles in each neural population. In **A** and **B**, dashed lines indicate the variances explained by PCA in each of the four neural populations without the shuffle.

Next, we examined whether the size of eigenvectors is different among the four neural populations, which represents the extent of neural modulation due to the probability and magnitude in each neural population as an arbitrary unit. The size of the eigenvectors was not significantly different (Figure 4D, left, PC1, Kruskal-Wallis test, n = 104, df = 3, H = 2.62, *P* = 0.45, right, PC2, n = 104, df = 3, H = 4.76, *P* = 0.19), but it strongly depended on a temporal resolution for the analysis (Figure S4). Size of the eigenvector decreased as the analysis window size decreased (Figure S4B, E, and F), although all results and conclusions described above were maintained across the window’s sizes (Figure S4A-D). The decrease in eigenvector size would be because signal-to-noise ratios generally decrease if the window size decreases. These effects were observed as the decrease in PCA performance (Figure S4A) and percentages of neural modulations in the conventional analyses (Figure S5).

Collectively, these observations suggested that the probability and magnitude of rewards could be detected and integrated within the activity of cOFC and VS neural populations as the expected value and risk-return signals in a stable state.

### Temporal structure of the neural population signals

To characterize temporal aspects of the VS and cOFC neural populations that yield the expected values signals, we compared temporal dynamics of all the four neural population signals. Specifically, we compared temporal patterns of the vector change exhibited by each neural population (Figure 6). At the time point after the cue onset when the monkeys initiated the expected-value computation, all the four neural populations developed eigenvectors (Figure 6A). Size of the eigenvectors increased and then decreased within a second, however temporal patterns of this change in size were different among the four neural populations. The onset latencies, detected by comparing to the vector size during the baseline period, seemed to be coincident for the cOFC, VS, and DS population, which was followed by a late noisy signal in the mOFC (Figure 6B). In contrast, the detected peak of vector size for each neural population seemed to appear at different times. To statistically examine these temporal dynamics at the population level, we used a bootstrap re-sampling technique (see Methods).

**Figure 6.**
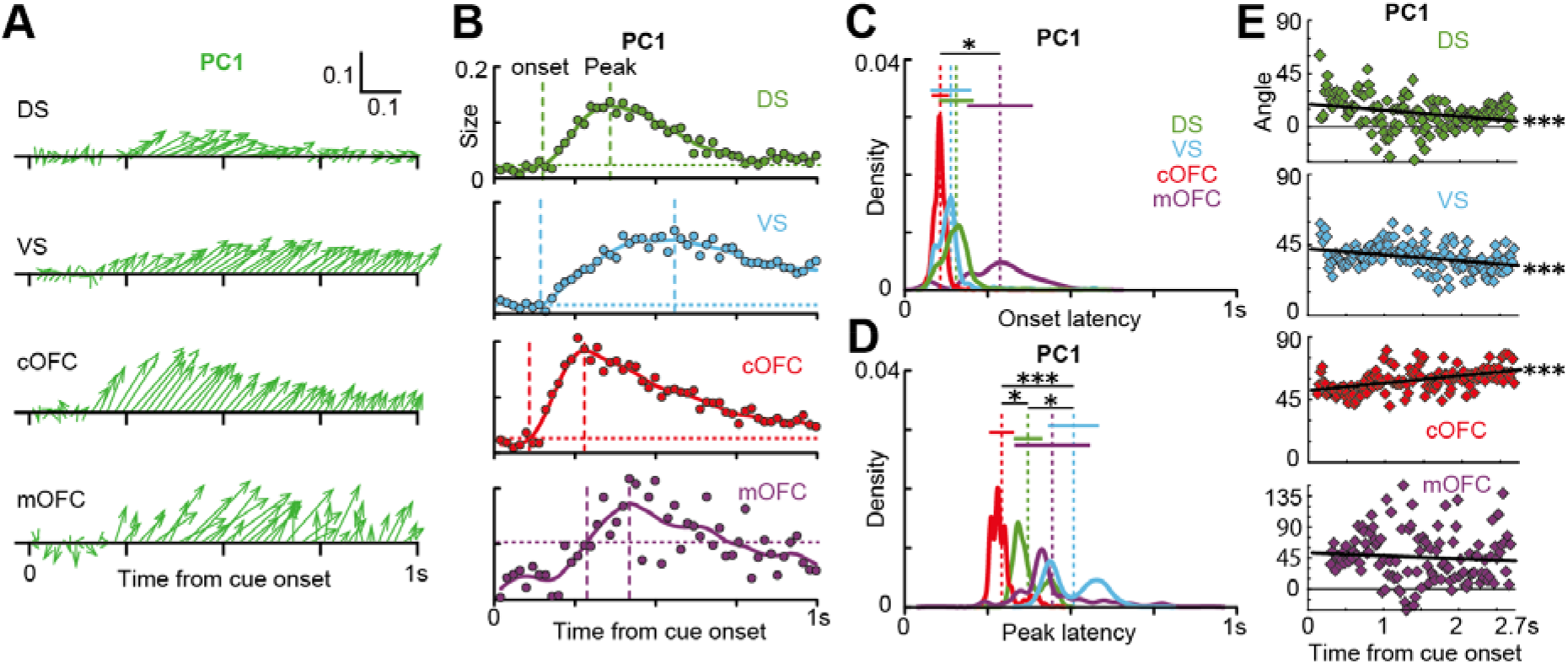
Gradual and sharp evolutions of neural population signals in the VS and cOFC. (**A**) Plots of the time series of eigenvectors for PC1 in 0.02 s analysis windows shown in a sequential order during 1 s after the cue onset. Horizontal and vertical scale bars indicate the eigenvectors for the probability and magnitude in arbitrary units, respectively. (**B**) Plots of the time series of vector size during 1 s after the cue onset. Horizontal dashed lines indicate three standard deviations of the mean vector size during the baseline period, a 0.3 s time period before the cue onset. Solid colored lines indicate interpolated lines using a cubic spline function to provide a resolution of 0.005 s. Vertical dashed lines indicate the onset (left) and peak (right) latencies for the changes in vector sizes. (**C**) Probability densities of the onset latencies of the four neural population signals. The probability densities were estimated using bootstrap re-samplings. Vertical dashed lines indicate the means. Horizontal solid lines indicate the bootstrap standard errors. (**D**) same as **C**, but for the peak latencies of the four neural population signals. (**E**) Plots of the time series of vector angle from the detected onset to the onset of outcome feedback. Solid black lines indicate regression slopes. In **C** and **D**, asterisks indicate statistical significance estimated using bootstrap re-samplings (*** and * indicate statistical significance at *P* < 0.001 and *P* < 0.05, respectively). In **E**, triple asterisks indicate a statistical significance of the regression slope at *P* < 0.001. Data for the PC2 is not shown.

The analysis revealed no significant difference in the onset latencies among cOFC, VS, and DS populations (Figure 6C, bootstrap re-sampling, onset latency, mean ± s.d., cOFC, 107.1 ± 26.0 ms, VS, 138,7 ± 61.3 ms, DS, 155.0 ± 52.4 ms) while these signals were followed by a late noisy signal in the mOFC (mOFC, 287 ± 98.8 ms). In contrast, if we compared the peak latencies (Figure 6D), the cOFC exhibited the earliest peak (292 ± 37.5 ms), followed by the DS (371 ± 43.0 ms), mOFC (444 ± 113.5 ms), and VS (508 ± 76.7), which exhibited the latest peak. Thus, the expected-value signal sharply developed in the cOFC in contrast to the gradual development in the VS. Note that mOFC signals were very noisy as indicated by the large variation in vector size during the baseline period (Figure 6B, bottom, see horizontal line).

Lastly, we examined temporal changes in vector angle, which indicate how fast the stable expected value signals evoked in the cOFC and VS (Figure 6E). As seen in the time series of the vector angle after detected onsets, the signals carried by the VS and cOFC neural populations at the early time period were almost 45° (i.e., expected values), indicating that these two neural populations integrate the probability and magnitude information into the expected values just after the numerical symbol appeared (see intercepts of the regression lines). Moreover, these two expected value signals were not the same, but rather idiosyncratic in each neural population: a gradual and slight shift of the vector angle directed to 90° (i.e., magnitude, cOFC, Figure 6E, regression coefficient, r = 5.31, n = 129, t = 6.04, df = 126, P < 0.0001) or 0° (i.e., probability, Figure 6E, VS, r = −3.91, n = 127, t = −4.16, df = 124, P < 0.0001) was observed toward the end of cue presentation. Similar to the VS, the DS population showed the same direction of the angle shift (Figure 6E, DS, r = −5.38, n = 127, t = −3.31, df = 124, P = 0.001). In contrast, such significant shift of the vector angle was not observed in the mOFC population (r = −4.30, n = 120, t = −0.94, df = 117, P = 0.351). These results indicated that the neural populations in both the VS and cOFC integrate the probability and magnitude information into the expected values immediately after the cue presentation, whereas their temporal dynamics were idiosyncratic in each of the two stable population signals.

## Discussion

Extraction of neural population dynamics is a recently developing approach for understanding computational processes implemented in the domain of cognitive and motor processing (Chen & Stuphorn, 2015; Churchland et al., 2012; Mante et al., 2013; Murray et al., 2017; Takei et al., 2017). This approach provides a mechanistic structure of neural population signals regarding temporal aspects, such as oscillatory activities during reaching (Churchland et al., 2012), co-activation patterns of spinal neurons and muscles (Takei et al., 2017), and a dynamic unfolding of task-related activity for perceptual decisions (Mante et al., 2013). Here, we found that the VS and cOFC neural populations maintain the stable expected-value signals at a population level (Figure 4). This is the first mechanistic demonstration of expected value signals embedded in multiple neural populations when monkeys compute the expected values from the numerical symbols cueing the probability and magnitude of rewards. The temporal dynamics of these two stable neural populations are unique in the aspect of time constants (Figure 6B-D) and gradual shifts of their structures (Figure 6E). These results indicate that the cOFC and VS compute the expected values as distinct, partially overlapping processes. These expected-value signals could be followed by the transition of neural population signals from evaluation to choices during economic decision makings (Chen & Stuphorn, 2015; Gardner et al., 2019; Yoo & Hayden, 2020).

### Two idiosyncratic expected-value signals in the cOFC and VS

State space analysis can detect both stable (Murray et al., 2017) and flexible (Mante et al., 2013) neural signals at a population level. In the present study, expected-value signals observed in the VS and cOFC were similarly stable in terms of the fluctuation of vector angles but were significantly different in terms of the temporal aspects (Figure 6). These signal properties indicate that information processing in these two brain regions were not same. For example, the fast cOFC signal may reflect the calculation of expected values from the probability and magnitude symbols like mental arithmetic, while the slow VS signal may reflect the secondary process to maintain the calculated expected-value information. It is known that the fronto-striatal projection plays a large role in a variety of cognitive functions (Haber & Knutson, 2010). Since the cOFC project to the VS anatomically, these two processes must act cooperatively through the cortico-basal ganglia loop. Indeed, both population signals were similar in terms of the heterogeneous signals carried by each individual neuron (Figure 2J and K) throughout the task trial (Figure 2N and O). However, these two expected-value signals were unambiguously distinctive in terms of their time course (Figure 6B-D) and gradual shift (Figure 6E). Therefore, the cOFC and VS may compute the expected values within each of these cortical and striatal local circuits in a co-operative manner.

Our results were consistent with human imaging studies, in which activity in the VS and cOFC represents value-related signals (Noonan et al., 2017; J. O’Doherty et al., 2004; Yan et al., 2016), but were not consistent with the evidences that value signals exist in human ventromedial prefrontal cortex (vmPFC) (Levy & Glimcher, 2012; Tom et al., 2007), which includes mOFC. The reasons for why the mOFC showed very weak signals related to all aspects of the expected values in the present study are unclear (Figure 2L and 4B). One possibility of this inconsistency may come from the species differences between human and non-human primates in the orbitofrontal network (Wallis, 2011). The mOFC is a part of the vmPFC, but the comparability between human and macaque monkeys remains elusive. Another possibility is that the vmPFC is not involved in simple information processing, such as association between cues and outcomes, but is involved in more complicated behavioral context for making economic decisions (Yamada et al., 2018) and setting of mood (Ongur & Price, 2000).

### Fluctuating signals in the DS and mOFC

Fluctuating signals were observed in the DS and mOFC because of the instability or weakness of the signals (Figure 4). The mOFC signal would not be completely meaningless, since the PCA performance in mOFC population was better than that in shuffle controls (Figure 5). However, the signal carried by mOFC population was weak (Figure 2L), indicating that the eigenvector fluctuation in mOFC population reflects weak signal modulations by the probability and magnitude. In contrast, PCA performance in the fluctuated DS population was equivalent to that in the cOFC population (Figure 4A), where the stable expected-value signal appeared. Moreover, considerable modulation of the neural activity in DS was observed in the conventional analyses (Figure 2I and M). Thus, the fluctuating DS signal must reflect some functional role employed by the DS neural population in detecting and integrating the probability and magnitude information. The DS signal fluctuated with a significant shift from relatively close to the expected values to probability signal (Figure 6E, top). The consistent direction of the shift between VS and DS populations implies that striatal neural populations may prefer probabilistic phenomena (Ma & Jazayeri, 2014; Pouget et al., 2013), whereas cOFC neural population may prefer magnitude, that is continuous variable.

### The expected value signals and economic choices

Economic choices employed in the brain seems to be dynamic sequential processes, such as the expected value computation, their comparison among alternatives, and making a choice. Recent findings suggest that these may or may not be discrete/continuous and overlapping processes until choices are made by the subject (Chen & Stuphorn, 2015; Yoo & Hayden, 2020). Neural correlates of the probability, magnitude, and the expected values have been extensively reported in a diverse set of brain regions (J. P. O’Doherty, 2014) without their dynamics. They may support a possibility that the expected value signals were computed in wide brain regions as a network as in the present study, although those signals were represented for the economic choice (Enomoto et al., 2020) or an array of alternative non-value related processes (J. P. O’Doherty, 2014), just as motor response for example. Although signals in the cOFC and VS were fluctuated (Figure 4B), these populations also signals expected values at the beginning of cue presentation (Figure 6A and E), suggesting the widespread evolution of the expected value signals might occur at very early time period through a reward circuitry.

### Significance of the population signals revealed by our state space analysis

State space analysis reveals temporal structures of neural population in multi-dimensional space for both cognitive (Murray et al., 2017) and motor tasks (Churchland et al., 2012; Takei et al., 2017). However, the interpretation of the extracted population structure depends on the way to use this procedure (Elsayed & Cunningham, 2017). In the present study, we did not seek to determine the population structure as a trajectory in neural state space, as was performed in previous studies. Instead, we aimed to detect the main features underscoring the population structure in the space of the probability and magnitude that compose the expected values. For this purpose, stability of the regression subspace is critical. We elaborately projected the neural firing rates into the regression subspace by preparing a completely orthogonal data matrix in our task design. Thus, our state space analysis is informative on whether and how the expected-value signals are composed of the probability and magnitude moment-by-moment as the series of eigenvectors.

## Conclusions

A dynamic integrative process of probability and magnitude underlies computation of expected values in restricted brain regions, i.e., cOFC-VS, which could occur before subjects make economic choices. Since the computation of expected values are followed by the choice process during economic choices, single cue task enables us to extract the integrative process for expected value computation. The existence of the neural population signals for expected-values is consistent with the expected values theory, whereas the co-existence of risk-return signals may reflect a behavioral bias for risk-preferences, a phenomenon observed across species (Stephens & Krebs, 1986; Yamada, Tymula, et al., 2013). The sharp and slow evolutions of expected-value signals in the cOFC and VS, respectively, indicate that each brain region has a unique time constant in the expected-value computation. When the monkeys perceive the probability and magnitude from numerical symbols, learned expected-values may be computed and recalled through the OFC-striatum circuit (Hirokawa et al., 2019). Our results indicate that the expected-value signals as observed in population ensemble activities are compatible with the framework of dynamical systems (Churchland et al., 2012; Mante et al., 2013).

## Acknowledgments

The authors appreciate the valuable comments of Tomomichi Oya, Tomohiko Takei, Tsuyoshi Setogawa, Jun Kunimatsu, Masafumi Nejime, Narihisa Matsumoto, Hiroshi Abe, and Takashi Kawai. In addition, the authors thank Takashi Kawai, Ryo Tajiri, Yoshiko Yabana, and Yuki Suwa for their technical assistance. Monkey FU was provided by NBRP “Japanese Monkeys” through the National Bio Resource Project of the MEXT, Japan.

## Materials and Methods

### Subjects and experimental procedures

Two rhesus monkeys were used (*Macaca mulatta,* SUN, 7.1 kg, male; *Macaca fuscata*, FU, 6.7 kg, female). All experimental procedures were approved by the Animal Care and Use Committee of University of Tsukuba (protocol#. H30.336) and performed in compliance with the US Public Health Service’s Guide for the Care and Use of Laboratory Animals. Each animal was implanted with a head-restraint prosthesis. Eye movements were measured using a video camera system at 120 Hz. Visual stimuli were generated by a liquid-crystal display at 60 Hz that was placed 38 cm from the monkey’s face when seated. The subjects performed the cued lottery task five days a week. The subjects practiced the cued lottery task for 10 months, after which they became proficient at making choices of lottery options.

### Behavioral task

#### Cued lottery tasks

Animals performed one of two visually cued lottery tasks: *single cue task* or *choice task*. Activity of neurons were recorded only during the single cue task.

##### Single cue task

At the start of each trial, the monkeys had 2 s to align their gaze to within 3° of a 1° diameter grey central fixation target. After fixating for 1 s, an 8° pie-chart providing information about the probability and magnitude of rewards was presented for 2.5 s at the same location as the central fixation target. The pie-chart was then removed and 0.2 s later a 1 kHz and 0.1 kHz tone (0.15 s duration) indicated reward and no-reward outcomes, respectively. The high tone preceded a reward by 0.2 s. The low tone indicated that no reward would be delivered. The animals received a fluid reward, for which magnitude and probability were indicated by the green and blue pie-charts, respectively; otherwise, no-reward was delivered. An inter-trial interval of 4 to 6 s followed.

##### Choice task

At the start of each trial, the monkeys had 2 s to align their gaze to within 3° of a 1° diameter grey central fixation target. After fixating for 1 s, two peripheral 8° pie-charts providing information about the probability and magnitude of rewards for each of the two target options were presented for 2.5 s, 8° to the left and right of the central fixation location. Grey 1° choice targets appeared at these same locations. After a 0.5 s delay, the fixation target disappeared, cueing saccade initiation. The animals were free to choose for 2 s by shifting their gaze to either target, within 3° of the choice target. A 1 kHz and 0.1 kHz tone of 0.15 s duration indicated reward and no-reward outcomes, respectively. The animals received the fluid reward indicated by the green pie-chart of the chosen target, with the probability indicated by the blue pie-chart of the chosen target; otherwise, no-reward was delivered. An inter-trial interval of 4 to 6 s followed.

#### Pay-off and block structure

Green and blue pie-charts indicated reward magnitudes from 0.1 to 1.0 mL in 0.1 mL increments and reward probabilities from 0.1 to 1.0 in 0.1 increments, respectively (Table S1). A total of 100 pie-charts was used. In the single cue task, each of the 100 pie-charts was presented once in random order. In the choice task, two out of the 100 pie-charts were randomly allocated to the two options. During one session of electrophysiological recordings, 20 to 50 trial blocks of the choice task were interleaved with 100 to 120 trial blocks of the single cue task.

#### Calibration of reward supply system

The precise amount of liquid reward was controlled and delivered to the monkeys with a solenoid valve. An 18-gauge tube (0.9 mm inner diameter) was attached to the tip of the delivery tube to reduce the trial-by-trial variability of reward supply. The amount of reward in each payoff condition was calibrated by measuring the weight of water with 0.002 g precision (hence, 2μL) on a single trial basis. This calibration method was the same as that in a previous study (Yamada et al., 2018).

### Electrophysiological recordings

We used conventional techniques for recording single neuron activity from the DS, VS, cOFC, and mOFC. The monkeys were implanted with recording chambers (28 × 32 mm) targeting the OFC and striatum, centered 28 mm anterior to the stereotaxic coordinates. The locations of the chambers were verified using anatomical magnetic resonance imaging. In each recording session, a stainless-steel guide tube was placed within a 1-mm spacing grid, and a tungsten microelectrode (1-3 MΩ, FHC) was passed through the guide tube. For recording of neurons in the mOFC and cOFC, the electrode was lowered until it approximated the bottom of the brain after passing through the cingulate cortex or dorsolateral prefrontal cortex or between them, respectively. For recording of neurons in the DS, the electrode was lowered until low spontaneous activity was observed after passing through the cortex and white matter. For recording of neurons in the VS, the electrode was lowered further until it passed through the internal capsule. At the end of the VS recording session, the electrode was occasionally lowered close to the bottom of the brain to confirm the recording depth relative to the bottom. Electrophysiological signals were amplified, band pass filtered, and monitored. Single neuron activity was isolated on the basis of spike waveforms. We recorded from the four brain regions of single hemisphere of the two monkeys: 194 DS neurons (98 and 96 from monkeys SUN and FU, respectively), 144 VS neurons (89, SUN and 55, FU), 190 cOFC neurons (98, SUN and 92, FU), and 158 mOFC neurons (64, SUN and 94, FU). The activity of all single neurons was sampled when the activity of an isolated neuron demonstrated a good signal-to-noise ratio (>2.5). Blinding was not performed. The sample sizes required to detect effect sizes (number of recorded neurons, number of recorded trials in a single neuron, and number of monkeys) were estimated with reference to previous studies (Chen & Stuphorn, 2015; Yamada, Inokawa, et al., 2013; Yamada et al., 2018). Neural activity was recorded during 100-120 trials of the single cue task. Note that presumed projection neurons (PANs, phasically active neurons) (Yamada et al., 2016) were recorded from the DS and VS.

### Statistical analysis

For statistical analysis, we used the statistical software package R (http://www.r-project.org/). All statistical tests for behavioral and neural analyses were two-tailed.

#### Effects of units on statistical analysis

In the present study, we used two variables for analyses; probability and magnitude. We defined the probability of reward from 0.1 to 1.0, and the magnitude of reward from 0.1 to 1.0 mL. Under this definition of units, the effects of probability and magnitude on the data were equivalent. Thus, the data were not standardized in the analyses.

### Behavioral analysis

We examined whether the monkey’s choice behavior depended on the expected values of the two options located left and right side to the center. We pooled choice data across all recording sessions, yielding 44,883 and 19,292 choice trials for monkeys SUN and FU, respectively. The percentage of right target choices was estimated for all combinations of the expected values of left and right target options.

#### Model Fitting

The percentage of choosing the right-side option was analyzed using a general linear model with binominal distribution,

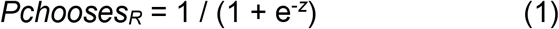

where the relation between *Pchooses_R_* and *Z* was given by the logistic function in each of the following three models: number of pies displayed (M1), probability and magnitude (M2), and the expected values (M3).

The first model, M1, assumed that the monkeys chose a target by comparing the number of pies for two targets,

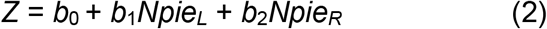

where *b*_0_ is the intercept, and *Npie_L_* and *Npie_R_* are the number of pies contained in left and right pie-chart stimuli, respectively. Values of *b*_0_ to *b*_2_ were free parameters and estimated by maximizing log likelihood.

The second model, M2, assumed that the monkeys chose a target by comparing the probability and magnitude of two targets,

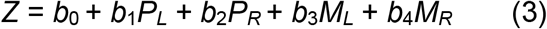

where *b*_0_ is the intercept; *P*_*L*_ and *P*_*R*_ are the probability of rewards for left and right pie-chart stimuli, respectively; and *M*_*L*_ and *M*_*R*_ are the magnitude of rewards for left and right pie-chart stimuli, respectively. Values of *b*_0_ to *b*_4_ were free parameters and estimated by maximizing log likelihood.

The third model, M3, assumed that the monkeys chose a target by comparing the expected values of rewards for two targets,

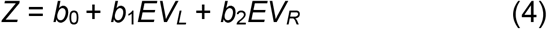

where *b*_0_ is the intercept, *EV_L_* and *EV_R_* are the expected values of rewards as probability times magnitude for left and right pie-chart stimuli, respectively. Values of *b*_0_ to *b*_2_ were free parameters and estimated by maximizing log likelihood.

#### Model comparisons

To identify the best structural model to describe the monkeys’ behavior, we compared the three models described above. In each model, we estimated a combination of the best-fit parameters to explain monkeys’ choice behavior. We compared their goodness-of-fits based on Akaike’s Information Criterions (AIC) and Bayesian Information Criterion (BIC) (Burnham & Anderson, 2004),

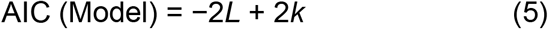

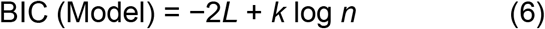

where *L* is the maximum log-likelihood of the model, *k* is the number of free parameters, and *n* is the sample size. After estimating the best-fit parameters in each model, we selected one model that exhibited the smallest AIC and BIC. To evaluate the model fits, we estimated a McFadden’s pseudo r-squared statistic using the following equation:

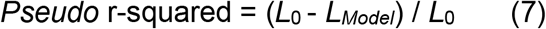

where *L_Model_* is the maximum log likelihood for the model given the data, and *L_0_* is the log likelihood under the assumption that all free parameters were zero in the model.

### Neural analysis

#### Basic firing properties

Peri-stimulus time histograms (PSTH) were drawn for each single neuron activity aligned at the visual cue onset. To display the histogram of neurons, the peak activity (maximum firing rate in each histogram) was detected for each neuron. The average activity curves were smoothed using a 50 ms Gaussian kernel (σ = 50 ms) and normalized by the peak firing rates. The percentage of neurons that showed an activity peak during cue presentation was compared among the four brain regions using a chi-squared test at *P* < 0.05. Basic firing properties, such as peak firing rates, latency of the peak, and duration of the peak activity (half peak width) were compared among the four brain regions using parametric and non-parametric tests, respectively, with a statistical significance level of *P* < 0.05. Baseline firing rates during 1 s before the appearance of central fixation targets were also compared with a statistical significance level of *P* < 0.05.

#### Estimation of neural firing rates through task trials

We analyzed neural activity during a 2.7 s time period from the onset of pie-chart stimuli to the onset of outcome feedbacks during the single cue task. To obtain a time series of neural firing rates through a trial, we estimated the firing rates of each neuron for every 0.1, 0.05, or 0.02 s time window (without overlap) during the 2.7 s period. No gaussian kernel was used.

#### Estimation of neural firing rates in a fixed time window

We analyzed neural activity during a 1 s time window after the onset of pie-chart stimuli during the single cue task. The 1 s activity was used for the conventional analyses below. No gaussian kernel was used.

### *Conventional analyses to detect neural modulations in each individual* neuron

#### Linear regression and model selection

For conventional and standard analyses of neural modulation by the values of probability and magnitude indicated by pie-chart stimuli, we used *linear regression* and *model selection analyses*. As mentioned above, we estimated the firing rate of each neuron during the 1 s time period after the onset of pie-chart stimulus during the single cue task. No gaussian kernel was used.

#### Linear regression

Neural discharge rates (*F*) were fitted by a linear combination of the following variables:

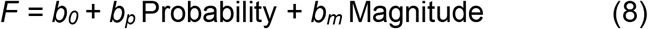

where Probability and Magnitude are the probability and magnitude of rewards indicated by the pie-chart, respectively. *b*_0_ is the intercept. If *b_p_* and *b_m_* were not 0 at *P* < 0.05, the discharge rates were regarded as being significantly modulated by that variable.

On the basis of the linear regression, activity modulation patterns were categorized into several types: “Probability” type with a significant *b*_p_ and without a significant *b*_m_; “Magnitude” type without a significant *b*_p_ and with a significant *b*_m_; “Expected value” type with significant *b*_p_ and *b*_m_ with both the same signs (i.e., positive *b*_p_ and positive *b*_m_ or negative *b*_p_ and negative *b*_m_); “Risk-Return” type with significant *b*_p_ and *b*_m_ with both having opposite signs (i.e., negative *b*_p_ and positive *b*_m_ or positive *b*_p_ and negative *b*_m_) and “non-modulated” type without significant *b*_p_ and *b*_m_. Note that risk-return types reflect high-risk high-return (prefer low probability and large magnitude) or low-risk low-return (prefer high probability and low magnitude).

#### Model selection

Neural discharge rates (*F*) were fitted by the following five models:

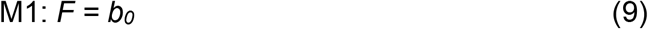

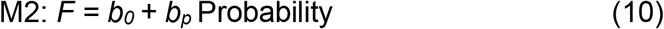

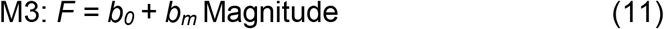

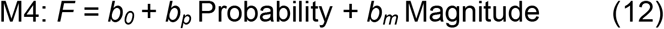

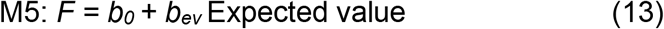

where Expected value is the expected value indicated by the visual pie-chart as the probability multiplied by magnitude (Table S1). *b*_0_ is the intercept. Probability and Magnitude are the probability and magnitude of reward indicated by the pie-chart, respectively. Among the five models, we selected one model that exhibited the smallest AIC or BIC.

If the selected model was M1, neurons were defined as the “non-modulated” type. If the selected model was M2, neurons were defined as the “Probability” type. If the selected model was M3, neurons were defined as the “Magnitude” type. If the selected model was M4 with the same signs of *b*_p_ and *b*_m_, neurons were defined as the “Expected value” type. If the selected model was M4 with opposite signs of *b*_p_ and *b*_m_, neurons were defined as the “Risk-Return” type. If the selected model was M5, neurons were defined as the “Expected value” type.

#### Application of conventional analyses to neural activity through task trials

We applied the three conventional analyses above (linear regression, AIC-based model selection, and BIC-based model selection) for the activity of neurons estimated at every time window in the four brain regions. As mentioned above, we estimated the firing rate of each neuron for every 0.1, 0.05, or 0.02 s time window (without overlap) during the 2.7 s period. No gaussian kernel was used. The activity modulation type was defined in each time window during the 2.7 s period. The analyses described percentages of the neural modulation types throughout cue presentation (Figure S5).

### Population dynamics using principal component analysis

#### Estimation of neuron firing rates through task trials

As mentioned above, we estimated the firing rate in each neuron every 0.1, 0.05, or 0.02 s time window (without overlap) during the 2.7 s period. No gaussian kernel was used.

#### Regression subspace

We used a linear regression to determine how the probability and magnitudes of rewards affected the activities of each neuron in the four neural populations. Each neural population was composed of all the recorded neurons in each brain region. We first prepared the probability and magnitude as 0.1 to 1.0 and 0.1 to 1.0 mL, respectively. We then described average firing rates of the neuron *i* at time *t* as a linear combination of the probability and magnitude in each neural population:

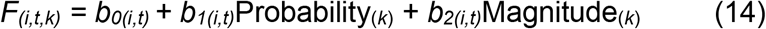

where *F_(i,t,k)_* is the average firing rates of neuron *i* at time t on trial *k,* Probability_(*k*)_ is the probability of rewards cued to the monkeys on trial *k;* Magnitude_(*k*)_ *is* the magnitude of rewards cued to the monkeys on trial *k*. The regression coefficients *b_0(i,t)_* to *b_2(i,t)_* describe the degree to which the firing rates of neuron *i* depend on the mean firing rates (hence, firing rates independent of task variables), the probability of rewards, and the magnitude of rewards, respectively, at a given time *t* during the trials.

We used the regression coefficients described in Eq. 14, above, to identify how the dimensions of neural population signals were composed from the probability and magnitudes as aggregated properties of individual neural activity. This step corresponds to the fundamental conceptual step of viewing the regression coefficients as a temporal structure of neural modulations by the probability and magnitude at the population level. Our procedures are analogous to the state-space analysis performed by Mante et al (Mante et al., 2013), in which the regression coefficients were used to provide an axis (or dimension) of the interested variables in multi-dimensional state space obtained by principal component analysis (PCA). In the present study, our orthogonalized task design allowed us to project the neural firing rates into the regression subspace reliably. It must be noted that our analyses were not aimed at describing population dynamics of the neural signals as a trajectory in the multi-dimensional task space, which is the standard goal of state space analysis.

#### Principal component analysis

We used a PCA to identify dimensions of the neural population signal in the orthogonal spaces composed of the probability and magnitude of rewards in each of the four neural populations. In each neural population, we first prepared a two dimensional data matrix *X* of size *N_(neuron)_*×*N _(C_*_×*T)*_; the regression coefficient vectors, *b_1(i,t)_* and *b_2(i,t)_* in Eq.14, whose rows correspond to the total number of neurons in each neural population and whose columns correspond to *C,* the total number of conditions (i.e., two: probability and magnitude); and T, the total number of analysis windows (i.e., 2.7 s divided by the window size). The PCs of this data matrix are vectors *v_(a)_* of length *N_(neuron)_*, total number of recorded neurons; if *N _(C_*_×*T)*_ is larger than *N_(neuron)_*, otherwise the length is *N _(C_*_×*T)*_. The PCs were indexed from the principal component explaining the most variance to the least variance. The eigenvectors were obtained by prcomp () function in the R software. It must be noted that we did not perform de-noising in the PCA (Mante et al., 2013), since we did not aim to project firing rates into state space. Instead, we intended to use the principal components to identify main features of the neural modulation signals at the population level through task trials.

#### Eigenvectors

When we applied the PCA to the data matrix *X*, we could deconstruct the set into eigenvectors and eigenvalues. The eigenvectors and eigenvalues exist as pairs with every eigenvector having a corresponding eigenvalue. In our analysis, the eigenvectors at time t represent a vector in the space of probability and magnitude. The eigenvalues at time t for the probability and magnitude are scalars, indicating how much variance is in the data in that vector. The first principal component is thus the eigenvectors with the highest eigenvalues. We analyzed eigenvectors for the first (PC1) and second principal components (PC2) in the following analyses. It must be noted here that we applied the PCA once in each neural population and thus the total variances contained in the data were different among the four populations.

#### Analysis of eigenvectors

We evaluated characteristics of the eigenvectors for PC1 and PC2 in each of the four neural populations in terms of the vector angle, size, and deviance in the space of probability and magnitude (Figure 3). The angle is the vector angle from the horizontal axis from 0° to 360°. The size is the length of the eigenvector. The deviance is the difference between vectors, and we estimated the deviance from the mean vector in each neural population. These three characteristics of the eigenvectors were compared among the four neural populations at *P* < 0.05 using the Kruskal-Wallis test and Wilcoxon rank-sum test with Bonferroni correction for multiple comparisons. The vector during the first 0.1 s was extracted from these analyses.

#### Shuffle control for PCA

To examine the significance of the population structure described by the PCA, we performed two shuffle controls. When we projected neural activity into the regression subspace, we randomized the data by shuffling in two ways. In the shuffled condition 1, *b*_*1(i,t)*_ and *b*_*2(i,t)*_ in Eq.14 were estimated with the randomly shuffled allocation of trial number *k* to the Probability_(*k*)_ and Magnitude_(*k*)_ only once for all time *t* in each neuron. This shuffle provided a data matrix *X* of size *N_(neuron)_*×*N _(C_*_×*T)*_, eliminating the modulation of probability and magnitude observed in the condition *C*, but retaining the temporal structure of these modulations across time. In the shuffled condition 2, *b_1(i,t)_* and *b_2(i,t)_* in Eq.14 were estimated with the randomly shuffled allocation of trial number *k* to the Probability_(*k*)_ and Magnitude_(*k*)_ at each time *t* in each neuron. This shuffle provided a data matrix *X* of size *N_(neuron)_*×*N _(C_*_×*T)*_, eliminating the structure across conditions and times. In these two shuffle controls, matrix *X* was estimated 1,000 times. The PCA performance was evaluated by constructing distributions of the explained variances for PC1 to PC4 (Figure 5). The statistical significance of the explained variances by PC1 and PC2 was estimated based on the bootstrap standard errors (i.e., standard deviation of the reconstructed distribution).

#### Bootstrap re-samplings for onset and peak latencies of the neural population signals

To detect onset and peak latencies of the population signal, we analyzed dynamic changes in population structure with the size of the eigenvectors in each of the four populations. We used a time-series of eigenvectors in 0.02 s analysis windows and estimated sizes of the time-series of vectors for PC1. To obtain smooth changes in vector size, a cubic spline function was applied with a resolution of 0.005 s (Figure 6B). Vector sizes during a baseline period were obtained by applying PCA to the matrix data *X* with time *t* from 0.3 s before cue onset to the onset of feedback (i.e., 3.0 s time period). A standard deviation of the vector sizes during the baseline period was obtained in each of the four neural populations. An onset latency of the population signal was defined as a time when the spline curve was above a value of more than three s.d. during the baseline period. Peak latency of the population signal was defined as the time from cue onset to the time when the maximum size of the vectors was obtained.

We estimated mean latencies of the onset and peak using a parametric bootstrap re-sampling method. In each neural population, the neurons were randomly re-sampled with a duplicate, and a data matrix *X* of size N_*(neuron)*_×N _*(C*×*T)*_ was obtained. The PCA was applied to the data matrix *X.* The time-series of the eigenvectors was obtained, and their sizes were estimated. The onset and peak latencies were estimated as above. This resampling was conducted 1,000 times, and distributions of the onset and peak latencies were obtained (Figure 6C and D). The statistical significance of the onset and peak latencies was estimated based on the bootstrap standard errors (i.e., standard deviation of the reconstructed distribution) (Efron. & Tibshirani., 1993).

## Supplemental Information

### Supplemental Figures

**Figure S1.**
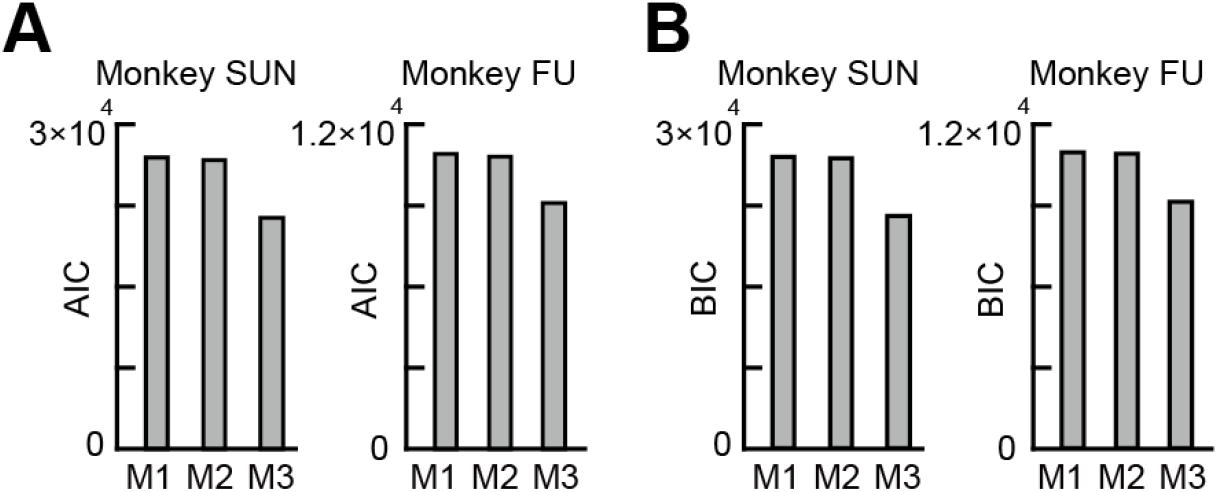
Model fits to the choice behavior of two monkeys. (**A**) Plots of the AIC values in monkey SUN and FU estimated in the three behavioral models, M1: number of pie. M2: probability and magnitude. M3: the expected values. (**B**) Same as **A**, but for the BIC values.

**Figure S2.**
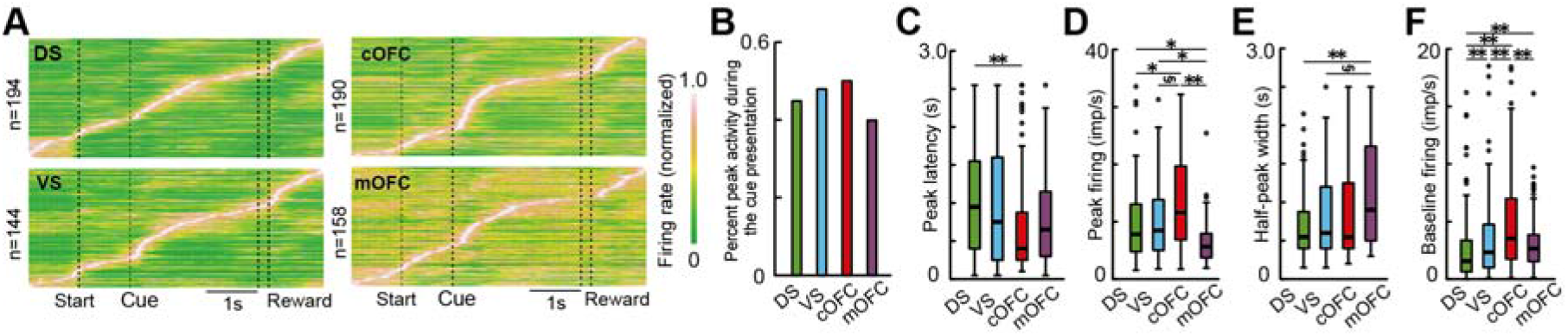
Basic firing properties of neurons in the DS, VS, cOFC, and mOFC. (**A**) Color map histograms of neuronal activities recorded from the four brain regions. Each horizontal line indicates the neural activity aligned to the cue onset, that was averaged for all lottery conditions. Firing rates of a neuron were normalized to the peak activity. (**B**) The percentages of neurons that showed an activity peak during the cue presentation. (**C**) Latency plots of the peak activity after the cue presentation. (**D**) Firing rates of the peak activity observed during the cue presentation. (**E**) Plots of the half-peak width, which indicates the phasic nature of activity changes. (**F**) Plots of the baseline firing rates during 1 s time period before the onset of the central fixation target. In **C-F**, asterisks indicate statistical significance among two neural populations using Wilcoxon rank-sum test with Bonferroni correction for multiple comparisons (**,*, and § indicate statistical significance at *P* < 0.01, *P* < 0.05, and 0.05 < *P* < 0.06 (close to significance), respectively).

**Figure S3.**
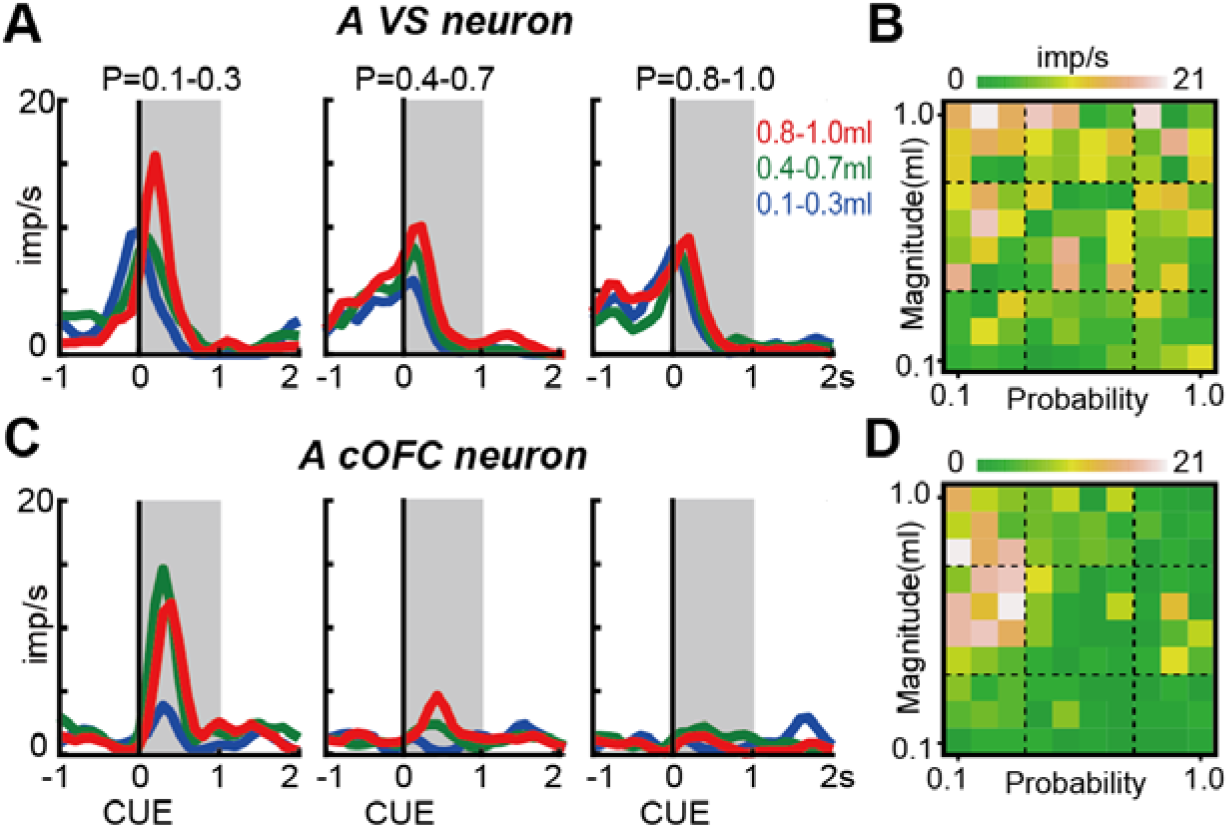
Risk-return signals detected by the conventional analyses. (**A**) An example activity histogram of a VS neuron modulated by both the probability and magnitude of reward with opposite signs (i.e., negative *b*_p_ and positive *b*_m_). The activity aligned to cue onset is represented for three different levels of the probability (0.1-0.3, 0.4-0.7, 0.8-1.0) and magnitude (0.1-0.3 mL, 0.4-0.7 mL, 0.8-1.0 mL) of rewards. Gray hatched areas indicate a 1 s time window to estimate the neural firing rates shown in **B**. The neural modulation pattern was defined as the Risk-Return type based on the linear regression and AIC-based model selection, and as Magnitude type based on the BIC-based model selection. Regression coefficients were −2.44 (*P* = 0.039) and 4.86 (*P* < 0.001) for the probability and magnitude, respectively. (**B**) Activity plots of the VS neuron during the 1 s time window shown in **A** against the probability and magnitude of rewards. (**C-D**) same as **A-B**, but for a cOFC neuron. The neural modulation type was defined as the Risk-Return type based on all the three analyses. Regression coefficients were −6.65 (*P* < 0.001) and 3.82 (*P* < 0.001) for the probability and magnitude, respectively.

**Figure S4.**
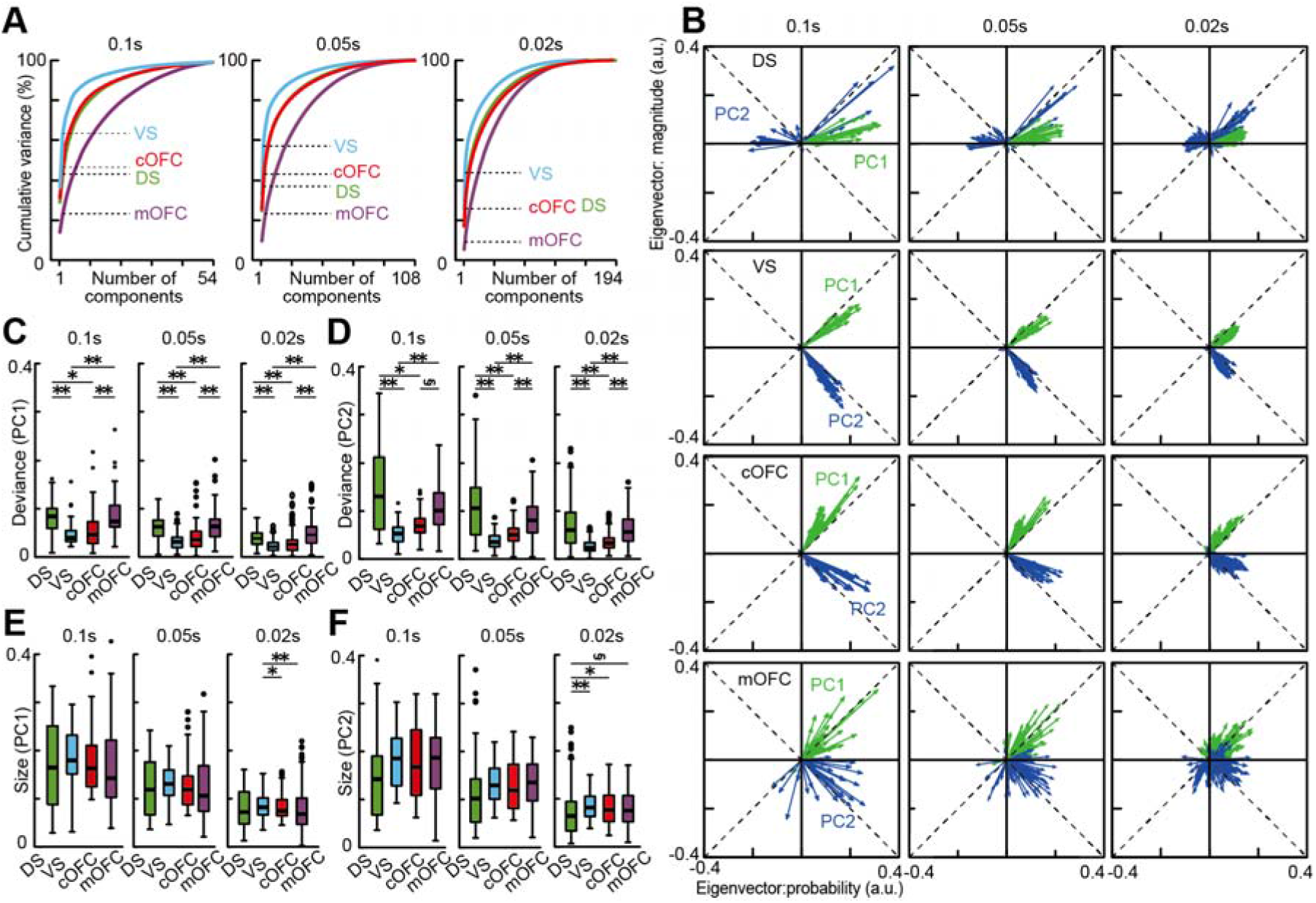
Effects of the analysis window size on the PCA. (**A**) Cumulative variances explained by PCA in the four neural populations. Dashed lines indicate the percentages of variance explained by PC1 and PC2 in each neural population. The size of the analysis windows was 0.1, 0.05, and 0.02 s, respectively. (**B**) Overlay plots of the series of eigenvectors in the four neural populations. The eigenvectors for PC1 and PC2 are shown. The size of the analysis windows was 0.1, 0.05, and 0.02 s, respectively. a.u. indicates arbitrary units. (**C**) Box plots of the deviance from the mean vector estimated in each neural population are shown for the PC1. (**D**) same as (**C**), but for the PC2. (**E**) Box plots of the vector size estimated in each neural population are shown for the PC1. (**F**) same as (**E**), but for the PC2. In **C-F**, asterisks indicate statistical significance between two neural populations using Wilcoxon rank-sum test with Bonferroni correction for multiple comparisons (**, *, and § indicate statistical significance at *P* < 0.01, *P* < 0.05, and 0.05 < *P* < 0.06 (close to significance), respectively).

**Figure S5.**
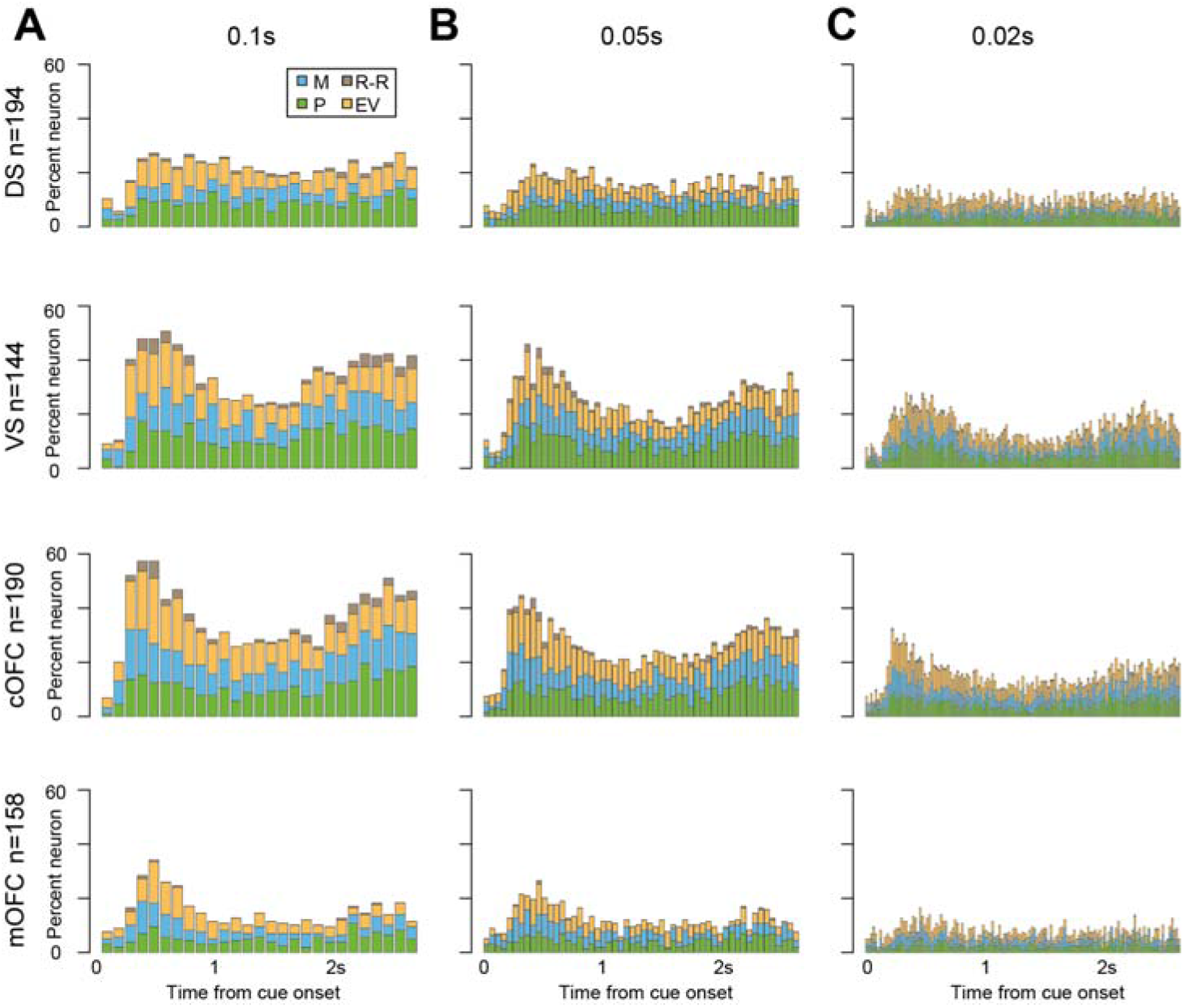
Effects of the analysis window size on the conventional analyses. (**A**) Percentage plots of the neural modulation type defined by using the model selection based on BIC values. The data for 0.1 s analysis windows is shown for the four neural populations. M: Magnitude type, P: Probability type, EV: Expected value type, R-R: Risk-Return type. (**B**) same as **A**, but for the data in 0.05 s analysis window. (**C**) same as **A**, but for the data in 0.02 s analysis window.

### Supplemental table

**Table S1.** Payoff matrix regarding probability and magnitude of rewards. Payoff matrix of the probability and magnitude of rewards indicated by the blue and green pie chart. The expected values [EV] defined as probability times magnitude are shown for the 100 possible combinations. All the 100 possible combinations of the reward probability and magnitude were used during the single cue and choice task.

| EV | Probability |  |  |  |  |  |  |  |  |  |
| --- | --- | --- | --- | --- | --- | --- | --- | --- | --- | --- |
|  | .10 | .20 | .30 | .40 | .50 | .60 | .70 | .80 | .90 | 1.0 |
| .10 | .01 | .02 | .03 | .04 | .05 | .06 | .07 | .08 | .09 | .10 |
| .20 | .02 | .04 | .06 | .08 | .10 | .12 | .14 | .16 | .18 | .20 |
| .30 | .03 | .06 | .09 | .12 | .15 | .18 | .21 | .24 | .27 | .30 |
| .40 | .04 | .08 | .12 | .16 | .20 | .24 | .28 | .32 | .36 | .40 |
| .50 | .05 | .10 | .15 | .20 | .25 | .30 | .35 | .40 | .45 | .50 |
| .60 | .06 | .12 | .18 | .24 | .30 | .36 | .42 | .48 | .54 | .60 |
| .70 | .07 | .14 | .21 | .28 | .35 | .42 | .49 | .56 | .63 | .70 |
| .80 | .08 | .16 | .24 | .32 | .40 | .48 | .56 | .64 | .72 | .80 |
| .90 | .09 | .18 | .27 | .36 | .45 | .54 | .63 | .72 | .81 | .90 |
| 1.0 | .10 | .20 | .30 | .40 | .50 | .60 | .70 | .80 | .90 | 1.0 |

